# Inferring copy number variation from gene expression data: methods, comparisons, and applications to oncology

**DOI:** 10.1101/2021.10.18.463991

**Authors:** Joseph Boen, Joel P. Wagner, Noemi Di Nanni

## Abstract

Copy number variations (CNVs) are genomic events where the number of copies of a particular gene varies from cell to cell. Cancer cells are associated with somatic CNV changes resulting in gene amplifications and gene deletions. However, short of single-cell whole-genome sequencing, it is difficult to detect and quantify CNV events in single cells. In contrast, the rapid development of single-cell RNA sequencing (scRNA-seq) technologies has enabled easy acquisition of single-cell gene expression data. In this work, we employ three methods to infer CNV events from scRNA-seq data and provide a statistical comparison of the methods’ results. In addition, we combine the analysis of scRNA-seq and inferred CNV data to visualize and determine subpopulations and heterogeneity in tumor cell populations.

## INTRODUCTION

Single-cell RNA-seq (scRNA-seq) is a powerful molecular profiling tool used to measure gene expression in individual cells [1]. The development of high-throughput sequencing technologies has made it possible to quantify gene expression in thousands of single cells for less than $1 USD per cell [2]. However, in addition to understanding the transcriptomic landscape, researchers are also interested in profiling the genomic landscape of tumors as well. Among genomic variants, copy number variants (CNVs) are an important genetic driver of cancer [3]. CNVs are genomic events in which the number of copies of a particular gene varies from individual to individual, or even from cell to cell.

Although there are tools for identifying CNVs using bulk whole-genome sequencing data [4], it has been more challenging to determine CNVs in single cells. Single-cell DNA sequencing has shown promise, but remains more challenging and is less ubiquitous than scRNA-seq. In addition, it remains technically challenging to sequence both the genome and the transcriptome from the same single cell [5]. With these challenges in mind, there has been interest in inferring CNVs from RNA-seq alone. However, this approach faces difficulties due to non-uniform coverage and dynamic variations in gene expression. Nevertheless, in recent years there have been several methods attempting this task.

In this work, we compare three methods that broadly represent the different approaches to inferring single-cell CNV profiles directly from scRNA-seq data: InferCNV [6], CaSpER [7], and CopyKAT [8]. InferCNV uses smoothed averages over gene windows; CaSpER uses a multi-scale signal processing framework; and CopyKAT uses a Bayesian segmentation approach. We compare these methods on a publicly available glioblastoma scRNA-seq dataset and measure their performance under varying conditions of the input gene expression data and algorithms’ parameters. Lastly, we combine the analysis of scRNA-seq data and the CNV profiles inferred from the scRNA-seq data to determine subpopulations and visualize intratumor heterogeneity in glioblastoma.

## MATERIALS AND METHODS

### Dataset

We used a publicly available glioblastoma scRNA-seq dataset (GEO accession GSE57872 [9]) of 5496 genes measured in 430 glioblastoma cells isolated from 5 patients (MGH26, MGH28, MGH29, MGH30, and MGH31) sequenced using the SMART-seq protocol. In addition, 1026 normal brain tissue bulk RNA-seq profiles from the Genotype-Tissue Expression (GTEx) project were included as control non-malignant samples intended to mimic normal single cells [10]. The normal tissue profiles were from the cerebellum, caudate, cortex, basal ganglia, cerebellar hemisphere, frontal cortex, and the hippocampus. For the InferCNV and CopyKAT algorithms, the glioblastoma and GTEx data were both accessed in the file “glio.wGtexBrain.counts.matrix.gz” at https://github.com/broadinstitute/inferCNV_examples/tree/master/glioblastoma. For the CaSpER algorithm, the normalized glioblastoma expression data and B-allele frequencies were accessed in the file “scell_gbm.rda” at https://github.com/akdess/CaSpER/tree/master/data.

### Workflows

InferCNV uses a corrected moving average of gene expression data to determine CNV profiles. Genes are sorted by absolute genomic position. That is, they are first ordered by chromosome, and then by genomic start position within the chromosome. The algorithm’s authors reason that averaging out the expression of genomically adjacent genes removes gene-specific expression variability and yields profiles that reflect chromosomal copy number variations. To further refine the CNV profile of tumor cells, InferCNV constructs the CNV profile of a known normal sample, and then for each gene and each cell, the normal sample is subtracted from the tumor sample to determine the final tumor CNV profile.

CaSpER uses a multiscale signal processing framework that combines two streams of information: gene expression and B-allele frequencies (BAF), both generated from aligned RNA-seq reads (BAM file). CaSpER first smooths both the BAF and expression signals using recursive iterative median filtering. At each scale, a hidden Markov model is applied to the smoothed expression signal and outputs five CNV states: (1) homozygous deletion, (2) heterozygous deletion, (3) neutral, (4) one-copy amplification, and (5) multi-copy amplification. For labels in state 2 and 4, CaSpER further incorporates information from the BAF signal. If there is an accompanying shift in the BAF signal at the segment, states 2 and 4 are corrected to states 1 and 5, respectively. BAF shifts are calculated by thresholding the smoothed signal. These thresholds are in turn calculated using a Gaussian mixture model fit on the pooled samples from all samples across a segment. As a result of these thresholds and corrections, CaSpER outputs discrete CNV classes: amplification, neutral, or deletion.

CopyKAT combines a Bayesian approach with hierarchical clustering. Like InferCNV and CaSpER, CopyKAT sorts genes by absolute genomic position, and then smooths and stabilizes the expression levels. CopyKAT next attempts to predict which cells among the input cells are diploid using hierarchical clustering. CopyKAT pools cells into several small hierarchical clusters and uses a Gaussian mixture model to calculate the variance of each cluster. The cluster with the least variance is assumed to be diploid and is used as a predicted normal cell reference CNV profile. Using this normal reference, relative gene expression profiles are calculated for the predicted tumor cells. The next step is to accurately determine chromosome breakpoints where CNV profiles change. Since scRNA-seq coverage is not uniform, CopyKAT employs Bayesian methods to best estimate the breakpoints. Specifically, CopyKAT uses a Poisson-gamma model and Markov Chain Monte Carlo to generate posterior means per genomic window. Then, Kolmogorov-Smirnov tests are used to determine whether to combine or keep separate the different windows. The final copy number values for each window are then calculated as the posterior averages for all genes spanning across chromosome breakpoints in each cell.

### Inputs, Outputs, and Technical Considerations

All three methods operate on the assumption that gene expression is correlated with copy number variation. That is, genes that are highly expressed are likely associated with copy number amplifications and genes that are lowly expressed are likely associated with copy number deletions. The key task is then to remove the confounding effect of natural fluctuations in gene expression levels, so that the excess or relative gene expression can be directly attributed to copy number variation.

Following this line of reasoning, all methods require a gene expression matrix (UMI counts) as input. InferCNV does not require labeled normal cell samples to work, but if none are provided it will simply use the average of all cells as the reference copy number profile. CopyKAT can work with either labeled or unlabeled normal cells as reference. If normal cells are assumed present but not explicitly labeled, CopyKAT will attempt to predict which cells are normal using hierarchical clustering. CopyKAT assumes that tumor cells are highly aneuploid while normal cells are diploid. The CopyKAT authors acknowledge this presents a limitation that can lead to potential misclassifications if the data only has a few normal cells, or when the tumor cells are nearly diploid. CopyKAT also requires expression data from at least five genes on every chromosome, whereas InferCNV allows for incomplete expression data. In addition to a gene expression matrix, CaSpER requires aligned RNA-seq BAM files to compute the BAF signal (Table 1).

**Table 1.** Details of three methods compared for inferring copy number variation from single-cell RNA-seq data.

| Method | Model | Input | Output | Tool |
| --- | --- | --- | --- | --- |
| CaSpER (7) | Multiscale signaling processing framework | - UMI count<br>- Aligned RNA seq | - Cell by gene CNV matrix, categorical values: amplification, deletion, neutral | - R package<br>https://github.com/akdess/CaSpER |
| CopyKAT (8) | Bayesian segmentation approach | - UMI count<br>- Optimal cell annotation | - Cell by gene CNV matrix, continuous values<br>- Tumor/normal cell classification | - R package<br>https://github.com/navinlabcode/copykat |
| InferCNV (6) | Smoothed averages of gene windows | - UMI count<br>- Cell annotation | - Cell by gene CNV matrix, continuous values | - R package<br>https://github.com/broadinstitute/inferCNV/wiki |

CaSpER outputs discrete CNV classes (deletion, neutral, or amplification) while InferCNV and CopyKAT output continuous CNV values. CopyKAT’s CNV values are normalized between −1 and 1 for deletion and amplification, respectively, with no CNV change from the reference normal cells given as 0. Similarly, InferCNV’s outputs are normalized between 0 to 2 with no CNV change from the reference normal cells given as 1.

## RESULTS

### Experiments

We compared the output of the three algorithms when varying the number of normal samples used as reference and whether those normal samples were labeled as normal or not. We ran InferCNV and CopyKAT using scRNA-seq data from all 430 glioblastoma tumor cells combined with 0, 5, 50, 500, or all 1,026 normal brain bulk RNA-seq samples. With CopyKAT, we ran each of those conditions twice: once when explicitly labeling which samples were normal and once allowing CopyKAT to predict which of the samples were normal. Since CaSpER does not require any normal samples, CaSpER was only run once using data from all tumor cells.

### Comparison of Technical Performance

To compare the CNV profiles of each method as a function of varying normal reference data, we sorted the estimated copy numbers by absolute genomic position and averaged the data over all tumor cells and, separately, all labeled or predicted normal samples. For CopyKAT and InferCNV, we averaged the values as reported directly from the method. For CaSpER, we encoded deletion as −1, neutral as 0, and amplification as 1. Each method applied to a different input thus produced a vector of CNV values, for a total of 16 different vectors: five from InferCNV, five from CopyKAT without labeling the normal samples as normal (“unlabeled”), five from CopyKAT with labeling the normal samples as normal (“labeled”), and one from CaSpER. Each element in the vector of CNV values represents the average CNV profile of each gene over all the tumor cells. This averaging procedure allows us to compare CaSpER, which has categorical outputs, to InferCNV and CopyKAT, which have continuous CNV profiles. From these averages, we computed the pairwise Spearman correlation over all pairs of vectors (Figure 1). We chose to compute the Spearman correlation, which only measures monotonicity between different variables, due to the arbitrariness of the CNV scales computed by each method. As expected, the row and one of the columns of low correlation corresponds to InferCNV and CopyKAT without any normal references. The remaining columns of low correlation correspond to unlabeled CopyKAT when provided with many normal samples. Since these columns do not appear in the labeled CopyKAT results, we conclude that these effects are caused by a failure in automatic cell type classification.

**Figure 1.**
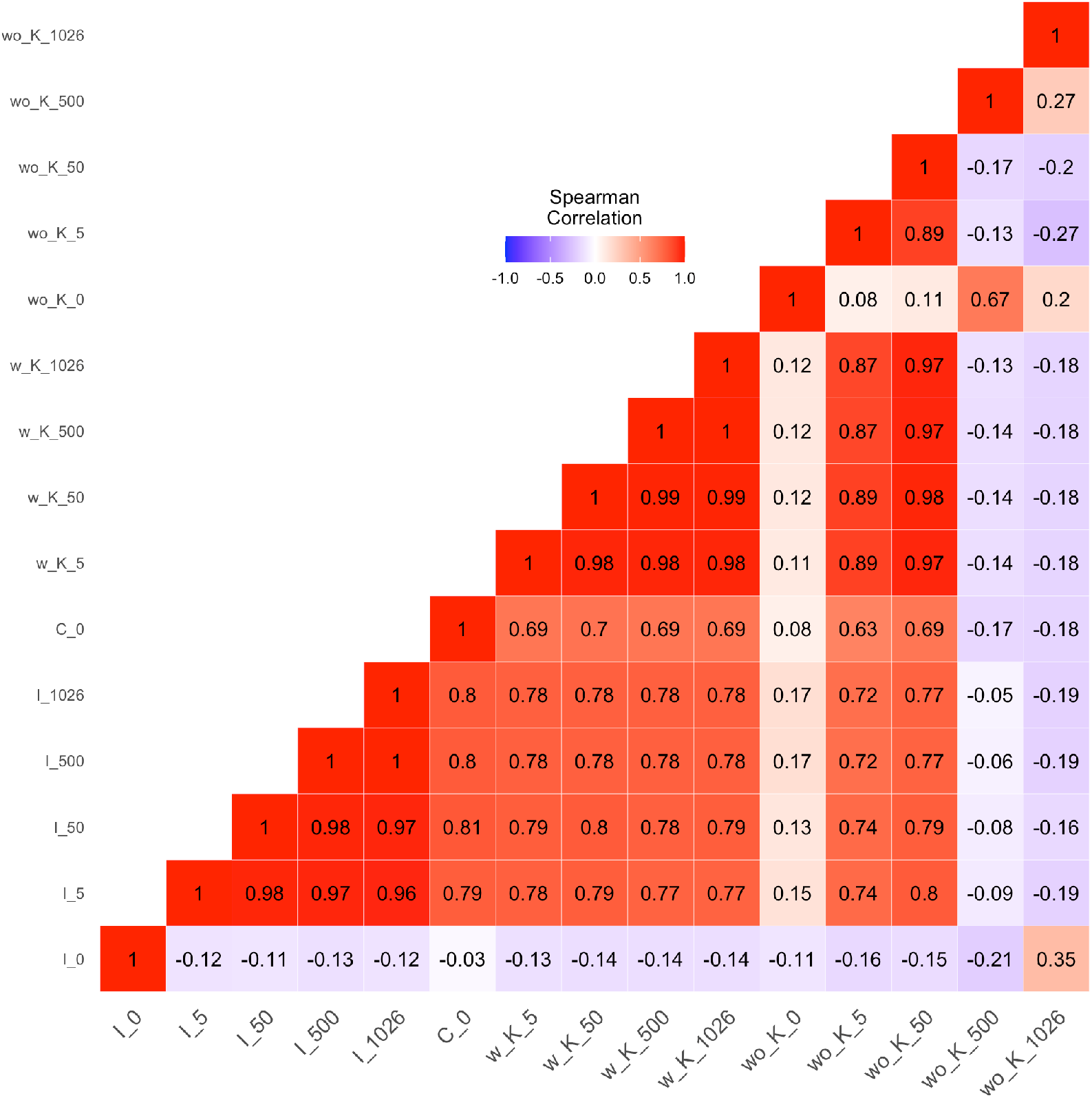
Inter-Methods Correlation. Spearman correlation matrix for averaged CNV profiles generated as described in the Experiments section. On the x-axis, from left to right, the columns indicate the results from InferCNV (I), CaSpER (C), CopyKAT provided with labeled normal samples (w_K), and CopyKAT provided with unlabeled normal samples (wo_K) for varying numbers of normal sample references (the number of normal samples used is given after the underscore).

In addition to showing the correlation plot, which shows the relations between all the methods and all the inputs, we highlight the performance of the three methods when using all tumor cells and all normal samples. The CNV profiles of only the tumor cells are shown across four approaches (Figure 2). Looking across the genes (columns), we can clearly see that distinct bands of CNV amplifications and deletions correspond to different patients (colorbars along rows), demonstrating inter-tumor heterogeneity. In contrast, there is little variation within a patient sample. Furthermore, CNV breakpoints are clearly present at the chromosome boundaries, as expected. Labeled CopyKAT, InferCNV, and CaSpER all produce similar CNV profiles. Interestingly, unlabeled CopyKAT appears to detect patient-specific CNV events but is unable to detect the population-wide amplification on chromosome 7 and deletion on chromosome 10.

**Figure 2.**
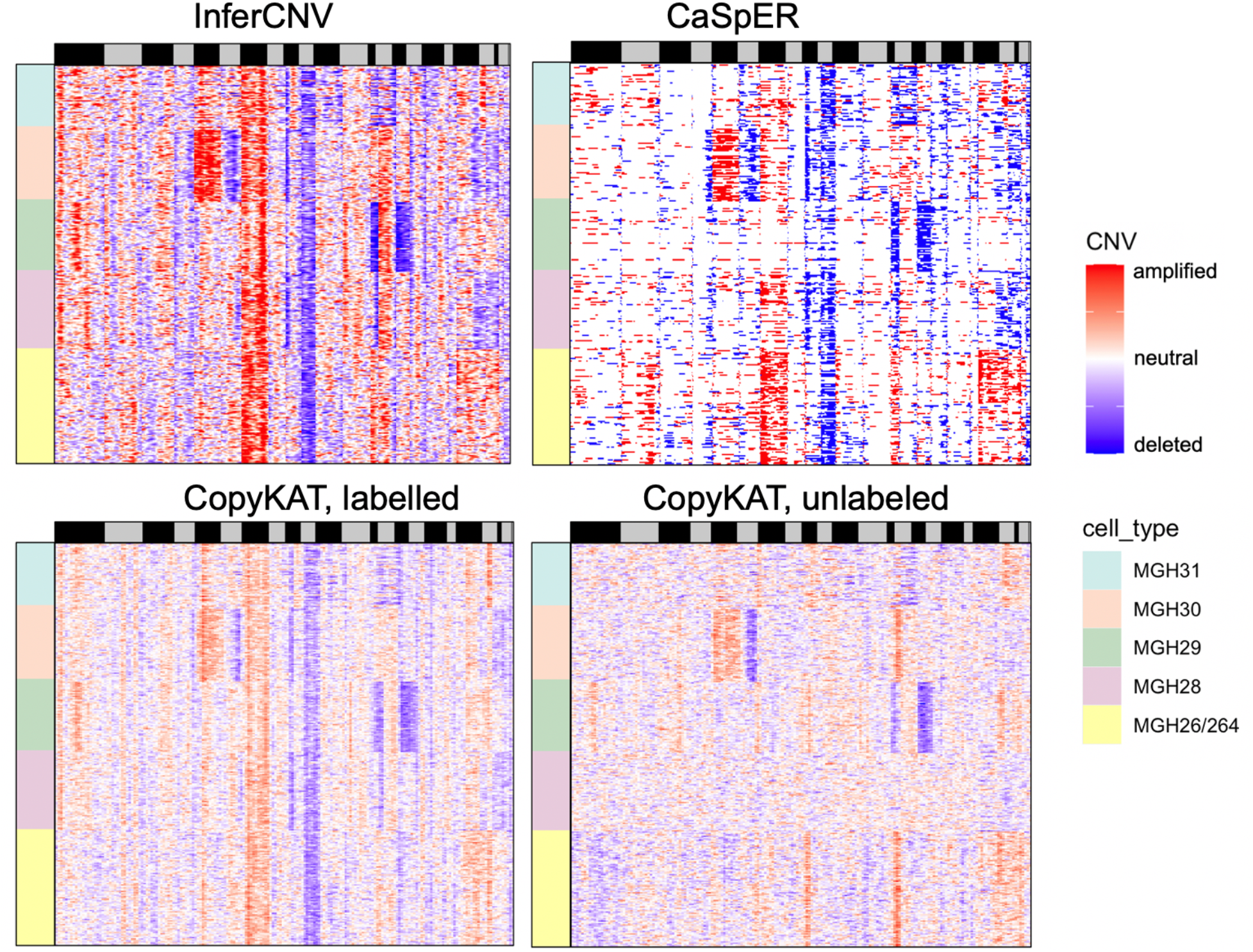
Copy Number profiles across methods. Copy number profiles generated by InferCNV and CopyKAT provided with 1026 normal samples, and CaSpER with 0 normal samples. CopyKAT is shown with and without cell labels. Rows correspond to individual cells across the five patient tumor samples (colorbars on the left) and columns correspond to genes ordered by chromosomal location (black and grey bars at the top indicate different chromosomes).

### Comparing gene expression and inferred CNV profiles

Tumor heterogeneity presents a major challenge in cancer diagnosis and treatment. Different tumor subtypes will lead to variable and distinct treatment responses and outcomes. In addition to inter-tumor variability, different cells within the same tumor might also harbor different mutations and exhibit different responses. Therefore, it is important to profile and resolve different subpopulations within a particular cancer, and then again within a particular patient. Clustering algorithms are commonly applied scRNA-seq data to detect subpopulations. From these clusters, one can apply differential gene expression analysis to determine biomarkers and cell types.

To this end, we wanted to compare cell clusters in the original gene expression data to those in the inferred CNV profiles. We applied principal component analysis and then generated uniform manifold approximation and projection (UMAP) plots for both the original gene expression data (Figure 3) and the corresponding inferred CNV profiles (Figure 4). Although we can clearly differentiate between bulk RNA-seq from normal brain samples and scRNA-seq from glioblastoma cells (Fig. 3A), the separation between different patients’ glioblastoma cells is less pronounced (Fig. 3B). When analyzing the inferred CNV profiles, we only considered the glioblastoma cells and combined results from the four algorithm conditions shown in Figure 3. In the UMAP projections of these inferred CNV profiles, the cells cluster first by patient (Fig. 4B) and then, within each patient, to some extent by algorithm (Fig. 4A). Thus, the inferred CNV values show more variability between patients than between methods. As expected by the heatmap visualizations, InferCNV, CaSpER, and annotated CopyKAT cluster very tightly. Surprisingly, un-annotated CopyKAT clusters closely as well. Lastly, the cells’ inferred CNV profiles (Fig. 4B) produce more distinct patient-specific clusters than the cells’ scRNA-seq data (Fig. 3B).

**Figure 3.**
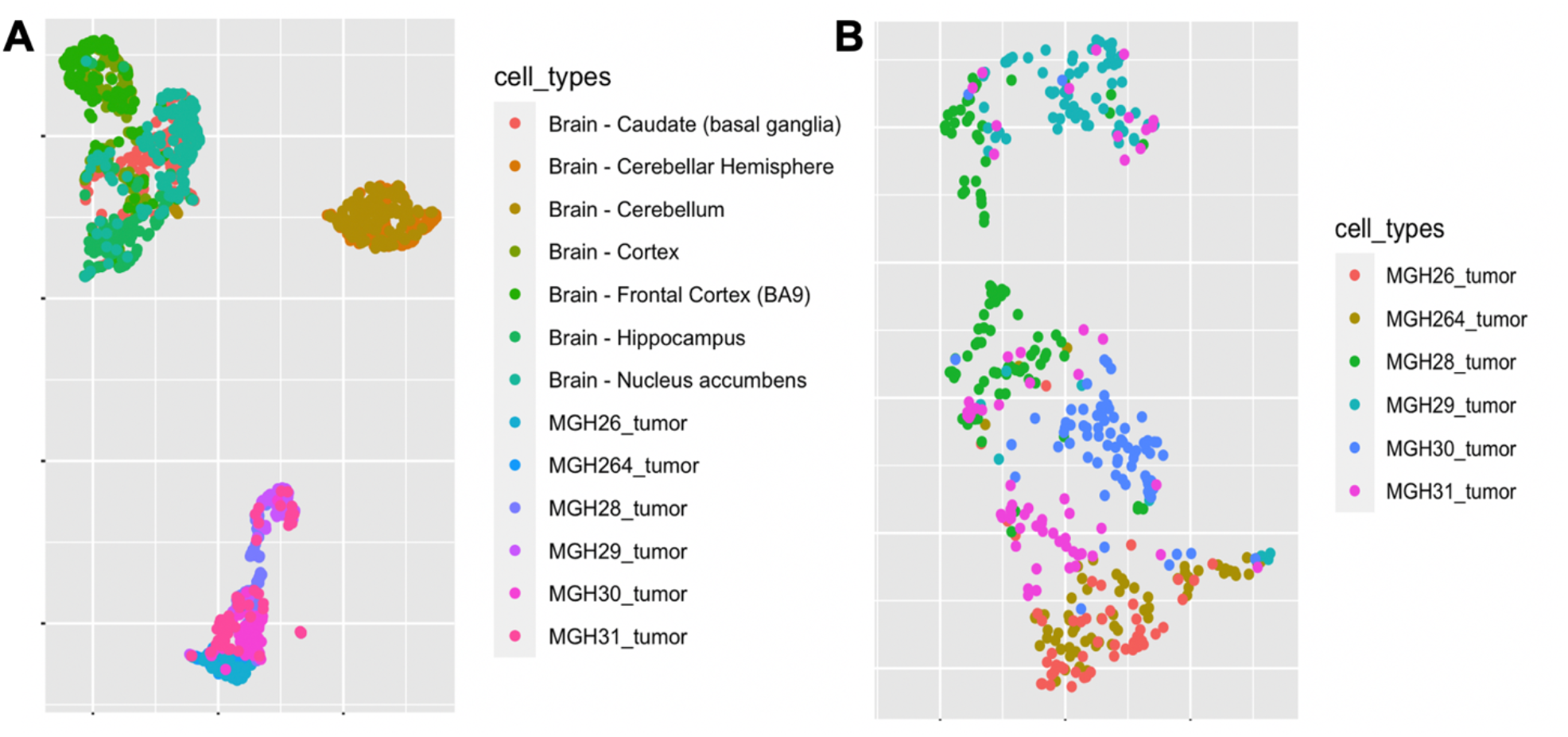
Gene Expression Clustering. UMAP projections of scRNA-seq data. Gene expression data is clustered using (A) tumor and normal cells, (B) only tumor cells. Gene expression data distinctly clusters tumor cells apart from normal brain tissue; however, within the tumor cells, the cells from different patients are not clearly separated.

**Figure 4.**
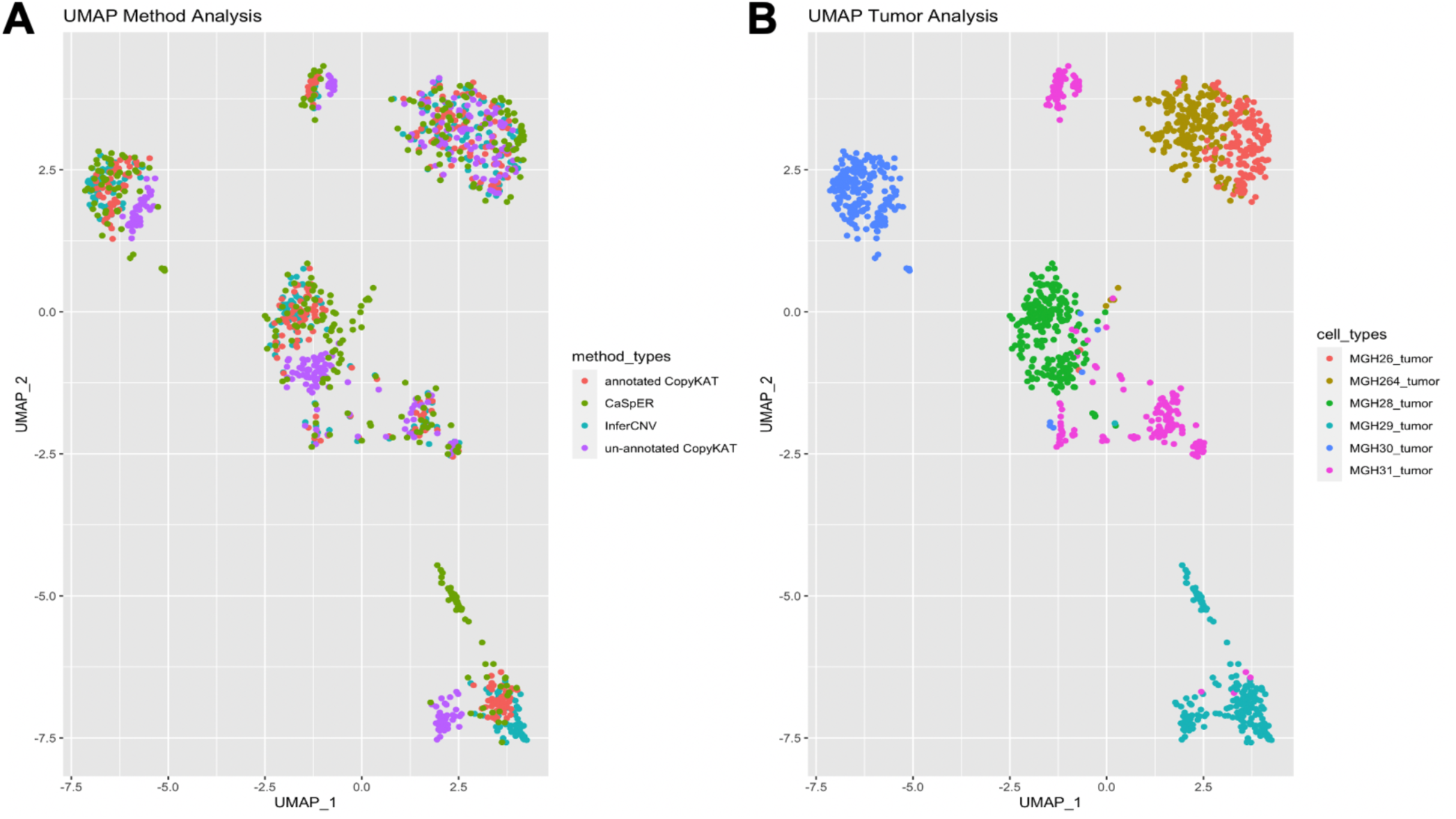
Copy Number Clustering. UMAP projections of inferred CNV data by method and by patient. The data shown in these figures correspond to the heatmaps shown in Figure 3 and were produced by the same experimental procedure. (A) The cells are colored by method. (B) The cells are colored by patient ID.

## DISCUSSION AND CONCLUSION

In this work we compared the results from three methods for inferring CNV profiles from gene expression data. We found that given sufficient reference information (e.g., BAM files, normal samples with annotations to use as reference profiles) all three methods were quite consistent in their predicted CNV profiles. However, in the absence of one or multiple pieces of reference information, performance varied wildly. Each method had their own limitations and strengths: InferCNV was a robust method, provided that at least some (as few as 5) normal samples were provided. CopyKAT was unable to differentiate between normal samples and tumor cells except in narrow circumstances with intermediate numbers of unlabeled normal samples relative to tumor cells, but performed as reliably as InferCNV when provided with normal sample annotations. By exploiting aligned RNA-seq information, CaSpER avoided the need for explicit normal references, but it only provided categorical CNV calls and not continuous CNV values like the other two methods.

Nevertheless, despite these caveats, the inferred CNV profiles proved to be a valuable tool for visualizing and determining tumor subpopulations and heterogeneity. Patient-specific distinctions that were obscured when solely analyzing gene expression data were more readily apparent when visualizing inferred CNVs. Interestingly, we discovered that inter-patient tumor CNV variability exceeded inter-method CNV variability, providing reassurance that despite the technical challenges of CNV estimation, the current methods provide effective approaches to uncovering inter-tumor heterogeneity.

Overall, we believe this work provides the most thorough comparison to date of methods for CNV inference from gene expression data and demonstrates the unique potential of CNVs for discovery in cancer biology.

## ACKNOWLEDGEMENTS

JB completed this work during the Novartis *A Summer of Science* Internship Program. The authors thank Sarah Szvetecz from the Oncology Data Science group at the Novartis Institutes for BioMedical Research (NIBR) for valuable discussions and support. And special thanks to Russette Lyons and Jessica Garver from NIBR for organizing the internship program.

